# Understanding the structural basis for differential binding of lapachol with PfHSP70s

**DOI:** 10.1101/2023.07.18.549604

**Authors:** Bikramjit Singh, Satinder Kaur, Komalpreet Kaur Sandhu, Chanchal Nainani, Vipul Upadhyay, Rachna Hora, Prakash Chandra Mishra

## Abstract

The 70 kDa heat shock proteins from *Plasmodium falciparum* (PfHSP70) are an important family of proteins that may be exploited for antimalarial design. Plant derived inhibitor ‘lapachol’ is reported to efficiently inhibit the exported PfHSP70-x while having poor activity against the parasite PfHSP70-1. In the present study, we have used *in silico* tools including molecular docking to understand the molecular basis for the above differences in lapachol inhibition activity against PfHSP70s. We found a significant gap in the binding energies and hence affinity of PfHSP70-1 and PfHSP70-x for lapachol with the latter having a better binding propensity. Our data highlight notable differences in the type of interactions between the two complexes. Detailed molecular analysis of the complexes has helped us to predict specific amino acid residues from both these PfHSP70 homologs that may be involved in lapachol binding. The above information may be utilized for design of PfHSP70 inhibitors with antimalarial potential.

## Introduction

Lapachol is a natural 1,4-napthoquinone (molecular formula C15H14O3) that was first derived from the bark of lapacho tree (*Tabebuiaavellanedae*) in 1882 [1]. It is substituted at positions 2 and 3 with hydroxy and 3-methylbut-2-en-1-yl groups respectively (Fig1a). While lapachol and its derivatives may be synthetically synthesized, many plants from diverse families like Bignoniaceae, Leguminosae, Malvaceae*etc*. are rich sources of this compound. Lapachol has vast pharmacological potential in cancer, sepsis and inflammatory diseases. It was found to be an effective cytotoxic agent against sixty cancerous cell lines in micromolar range. This phenolic metabolite is also reported to have diverse activities against bacteria, viruses, helminths, fungi, pests and protozoans (*Leishmania* and *Plasmodium*)[2][3]. Lapachol and its analogs displayed activity against leishmanial promastigotes (*L. amazonensis and L. braziliensis*) with IC_50_ values in the lower micromolar range [2]. Lapachol exhibited antimalarial activity in growth inhibition assays conducted on cultured *Plasmodium falciparum* (Pf) erythrocytes by Cockburn *et al*[4]with an IC_50_ value of 18.67µM. A lapachol derivative ‘Atovaquone’ has been approved to treat *Pneumocystis* caused pneumonia, toxoplasmosis and malaria [5].

Pf, the causative agent for most malaria associated fatalities overcomes several challenges posed by the intracellular host environment, particularly oxidative stress and temperature changes brought on by febrile episodes of malaria [4].Expression of a family of four heat shock proteins of approximately 70 kDa (PfHSP70) in Pf is a component of this adaptation. These are the well-studied PfHSP70-1 (nuclear & cytosolic), the endoplasmic reticulum (ER) expressed PfHSP70-2, the mitochondrial PfHSP70-3 and PfHSP70-x, which is exported to host erythrocyte cytoplasm [6]. HSP70s from humans and pathogens form an important group of drug targets that are being exploited for treatment of both infectious and non-infectious diseases owing to their critical roles in protein homeostasis. Therefore, PfHSP70s have been at the center stage for design of antimalarials by small molecule inhibitors. These include several classes of compounds like pyrimidinones, ATP analogues, spergualins, peptides *etc*. which are known to regulate HSP70 activity [7].

HSP70s are chaperones that function to prevent protein misfolding, assist refolding of misfolded proteins and prevent protein aggregation [1]. They interact with co-chaperones (PfHSP40s) via their N-terminal ATPase domain (also called nucleotide binding domain – NBD) and enhance binding of substrate proteins through their C-terminal substrate binding domain (SBD) [8]. Lapachol is reported to bind with the ATPase domains of PfHSP70-1 andPfHSP70-xto inhibit their aggregation suppression activity [3][4]. While lapachol inhibited both the basal and PfHSP40 stimulated ATPase activity of PfHSP70-x in a concentration dependent manner, no significant effect of this compound was seen on PfHSP70-1. We have therefore performed an *in silico* structural analysis of lapachol bound PfHSP70-1 and PfHSP70-x to understand differences in their response to the inhibitor.

## Materials and Methods

### Sequence and structure retrieval

The three dimensional structure of PfHsp70-x NBD (PlasmoDB ID: PF3D7_0831700; PDB ID: 6S02) was retrieved from PDB database [9], [10]. Since the structure of NBD of PfHsp70-1 (PlasmoDB ID: PF3D7_0818900) was not available on PDB, its sequence was retrieved from PlasmoDB to predict the structure of this protein domain. The structure of the Hsp70 inhibitor ‘lapachol’ was downloaded from PubChem (CID: 3884)[11].

### Molecular modeling

The structure of NBD of PfHsp70-1 was predicted using Swiss Model, a homology modeling server [12]. For this, the downloaded sequence of PfHsp70-1 was used to find suitable templates by performing blastp against PDB [13].The closest template was selected for molecular modeling based on percent identity and query coverage. The obtained model was validated for correctness by using Ramachandran plot and ERRAT [14].

### Molecular docking with lapachol and interaction analysis

Structures of PfHSP70-1, PfHSP70-x and lapachol were prepared for docking inChimera to generate PDBQT files [15]. Docking of lapachol was done with three dimensional structures of NBDs of both the proteins at a site adjacent to the nucleotide binding site (site reported to bind a small molecule inhibitor HEW) [16]. For this, grid boxes centered on the HEW binding pocket were used. Docking was done through AutoDockVina in Chimera leading to generation of multiple binding poses for each docking experiment [17]. The best poses were selected for each of the docked complexes and binding energies computed. Amongst these, the binding pose with best binding energy was analyzed to identify individual residues involved in binding using LigPlot+[18].

## Results and discussion

### Structures of PfHSP70-x, PfHSP70-1 and lapachol

The structure of lapachol (PubChem CID: 3884) was downloaded and visualized in Chimera (Fig 1a). Three-dimensional coordinates of NBD of PfHSP70-x (PlasmoDB ID: PF3D7_0831700) were obtained from PDB (PDB ID: 6S02) (Fig 1b). Since the structure of PfHSP70-1 (PlasmoDB ID: PF3D7_0818900) is not solved experimentally, we decided to predict the structure of PfHSP70-1 NBD using homology modeling. For this, Blastp was performed against PDB to identify the most suitable template. 6S02, i.e., ADP-bound NBD of PfHSP70-x was found to have maximum identity (85.09%) with PfHSP70-1 (57% query coverage). PfHSP70-1 and PfHSP70-x are also considered closest homologs in Pf as the full-length proteins share 73% sequence identity [20]. Selection of the template was followed by structure prediction using Swiss model (Fig 1c). The obtained structure was validated using Ramachandran plot analysis. None of the residues in the homology modeled structure were present in the disallowed region.

**Fig 1:**
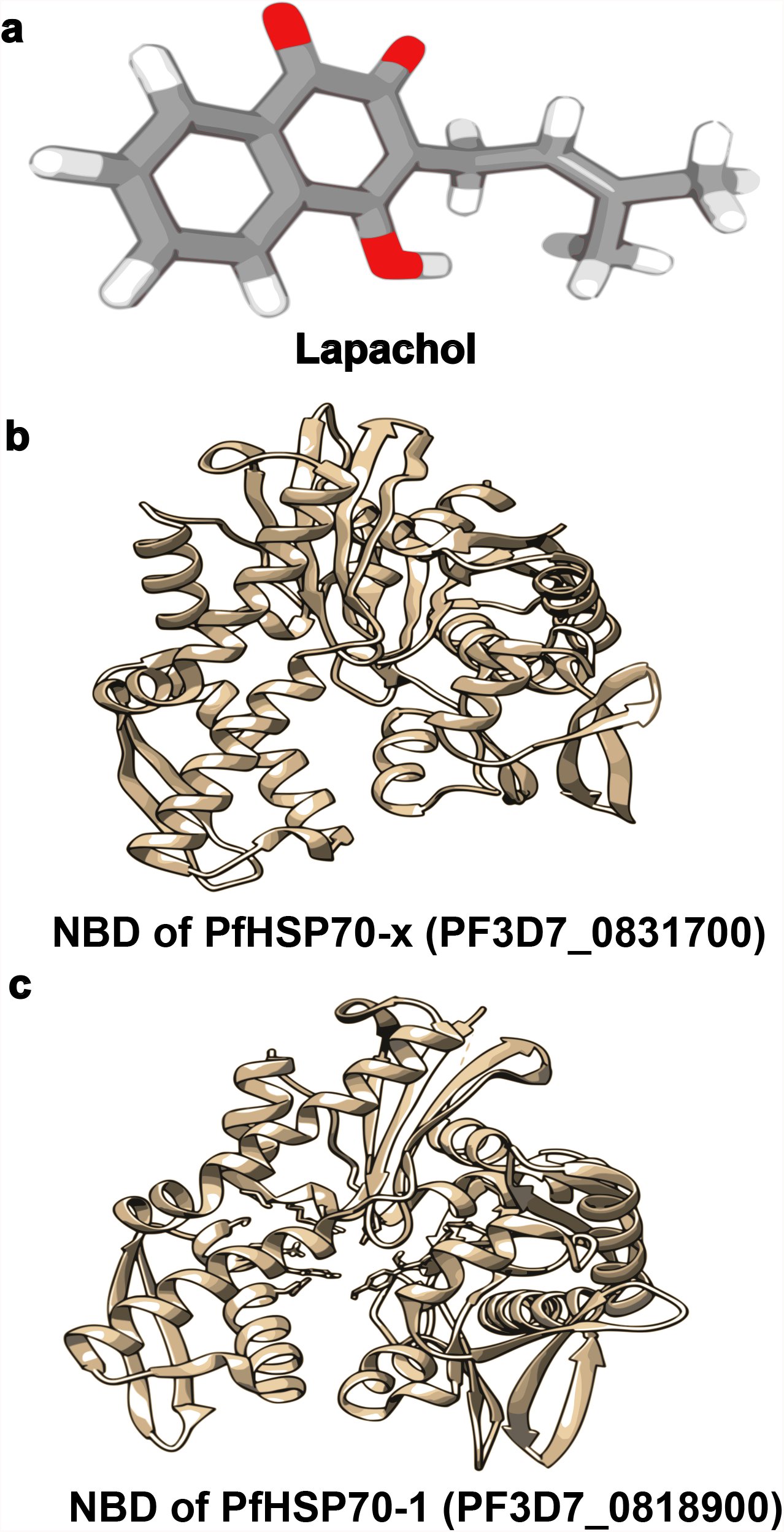
Molecular structures of proteins and ligand. a. Structure of lapachol downloaded from PubChem in SDF format (CID: 3884). b. Crystallographic structure of NBD of PfHSP70-x downloaded from PDB (6S02). c. Three-dimensional structure of NBD of PfHSP70-1 obtained from Swiss Model.

### Molecular docking and analysis

Mohamad *et al* had screened 233 chemical fragments for binding to the ATPase domain (NBD) of PfHSP70-x leading to identification of two binding sites on the molecule [16]. These were the nucleotide binding pocket and a site proximal to it, with most tested ligands including HEW binding at the proximal site. The amino acid residues of PfHSP70-x interacting with HEW were identified on the basis of crystallographic structure of PfHSP70-x-HEW complex (PDB ID 7OOG) (Table 1). Sequence alignment of PfHSP70-x and PfHSP70-1 helped us to mark the interacting residues on the latter (Table 1). We now selected grid boxes on PfHSP70-1 andPfHSP70-xcentered at the HEW binding site, and docked lapachol with coordinates of both the PfHSP70s (Fig 2). The docked complexes showed lapachol nestled in the HEW binding cavity on PfHSP70s (Fig 3).Our results revealed that lapachol bound more strongly to PfHSP70-x (binding energy: -8.774) as compared to PfHSP70-1(binding energy: -7.355). These values clearly indicate why lapachol efficiently inhibits the ATPase activity of PfHSP70-x, but has no significant effect on PfHSP70-1 action [3][4].

**Table 1:**
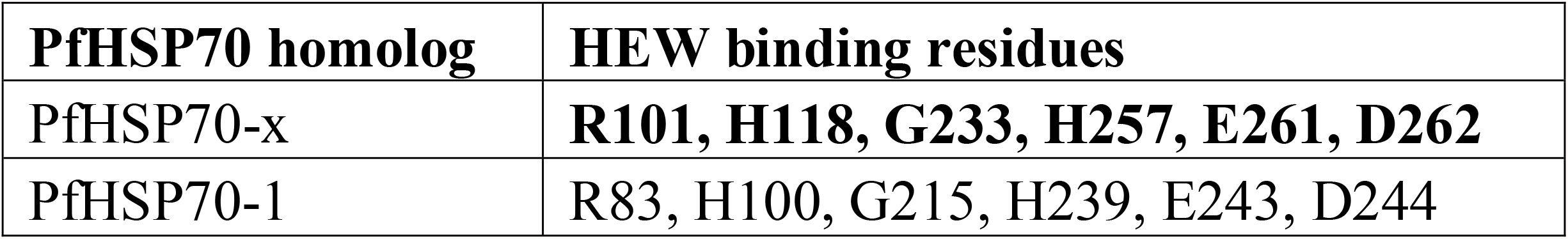
HEW binding sites on PfHSP70s. Amino acid residues of PfHSP70-x interacting with HEW identified on the basis of crystallographic structure of PfHSP70-x-HEW complex (PDB ID 7OOG) are shown in bold. PfHSP70-1 residues interacting with HEW determined by sequence alignment are tabulated.

| **PfHSP70 homolog** | **HEW binding residues** |
| --- | --- |
| PfHSP70-x | **R101, H118, G233, H257, E261, D262** |
| PfHSP70-1 | R83, H100, G215, H239, E243, D244 |

**Fig 2:**
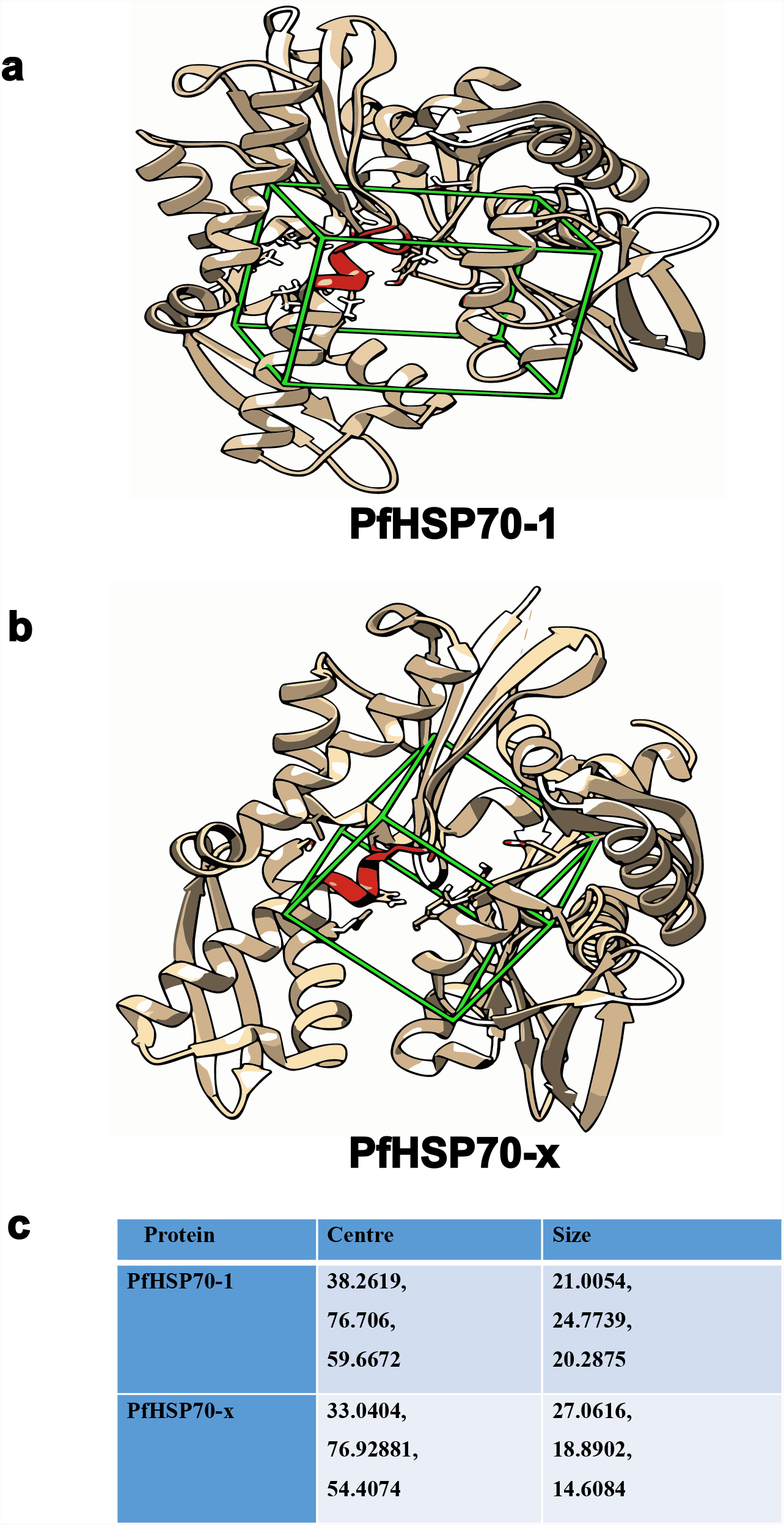
Grid boxes for molecular docking. Grid boxes (green) marked around HEW binding sites (red) on NBDs of a. PfHSP70-1 and b. PfHSP70-x. Protein molecules are shown as beige ribbon diagrams. c. Center coordinates and sizes of the selected grid boxes for HEW binding sites on PfHSP70 homologs are tabulated.

**Fig 3:**
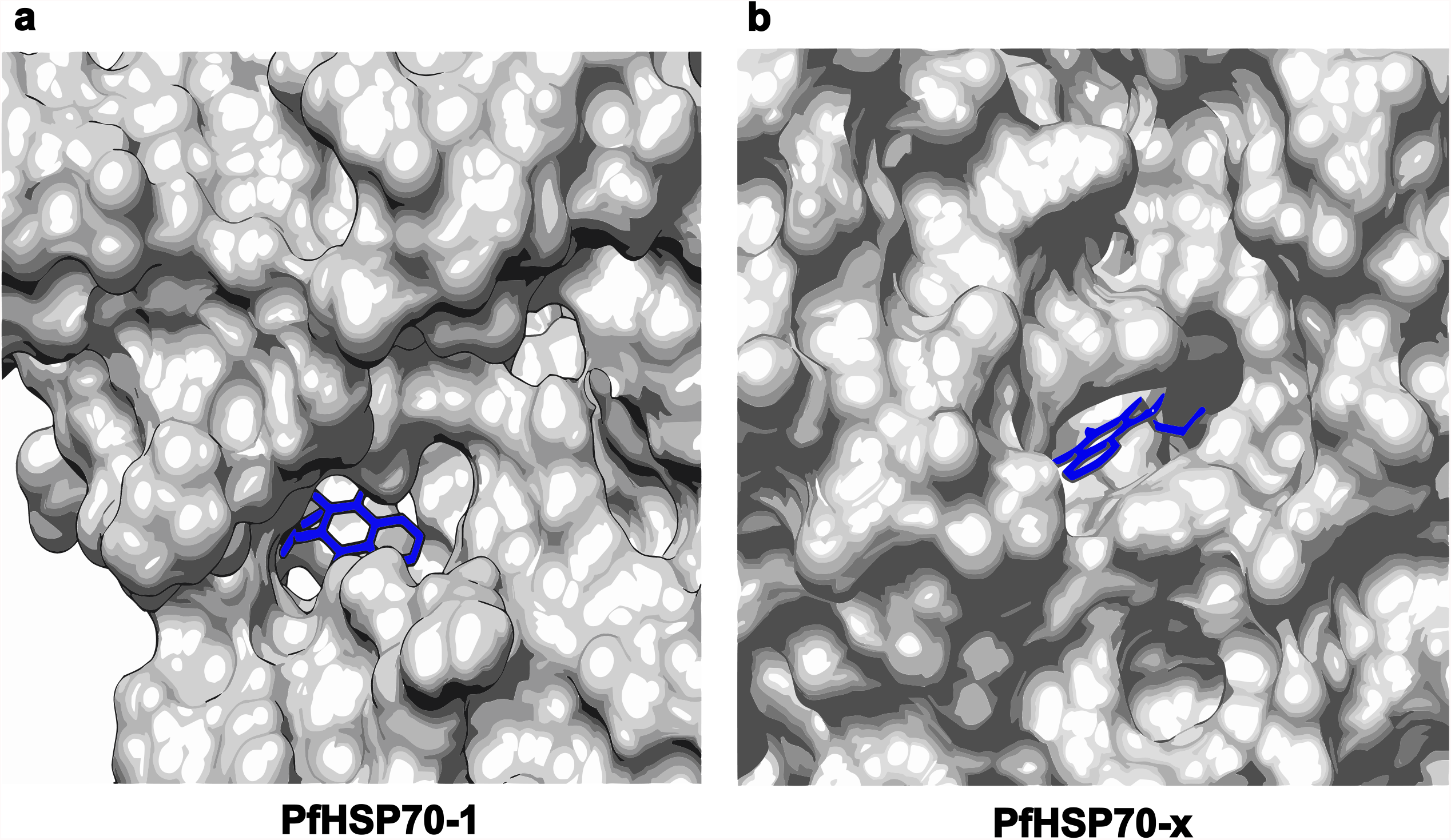
Molecular surface representation of PfHSP70-lapachol complexes. Lapachol (blue) docked on a. PfHSP70-1 and b. PfHSP70-x. Protein molecules are shown in grey.

The PfHSP70-lapachol complexes obtained from docking studies wereanalysed by using LigPlot+ to identify specific amino acid residues on the chaperone that are involved in inhibitor binding (Fig 4; Table 2). All the PfHSP70-1 contacts with lapacholand most from PfHSP70-x were non-bonded interactions like hydrophobic or Van der Waals. Interestingly, these interactions involved distinct amino acid residues from both proteins (Table 2). While hydrogen bonding also played a role in PfHSP70-x-lapachol binding, these contacts were absent from the PfHSP70-1-lapachol complex. Figure 5pictorially represents residues of the studied PfHSP70s involved in lapachol binding (Fig 5).

**Fig 4.**
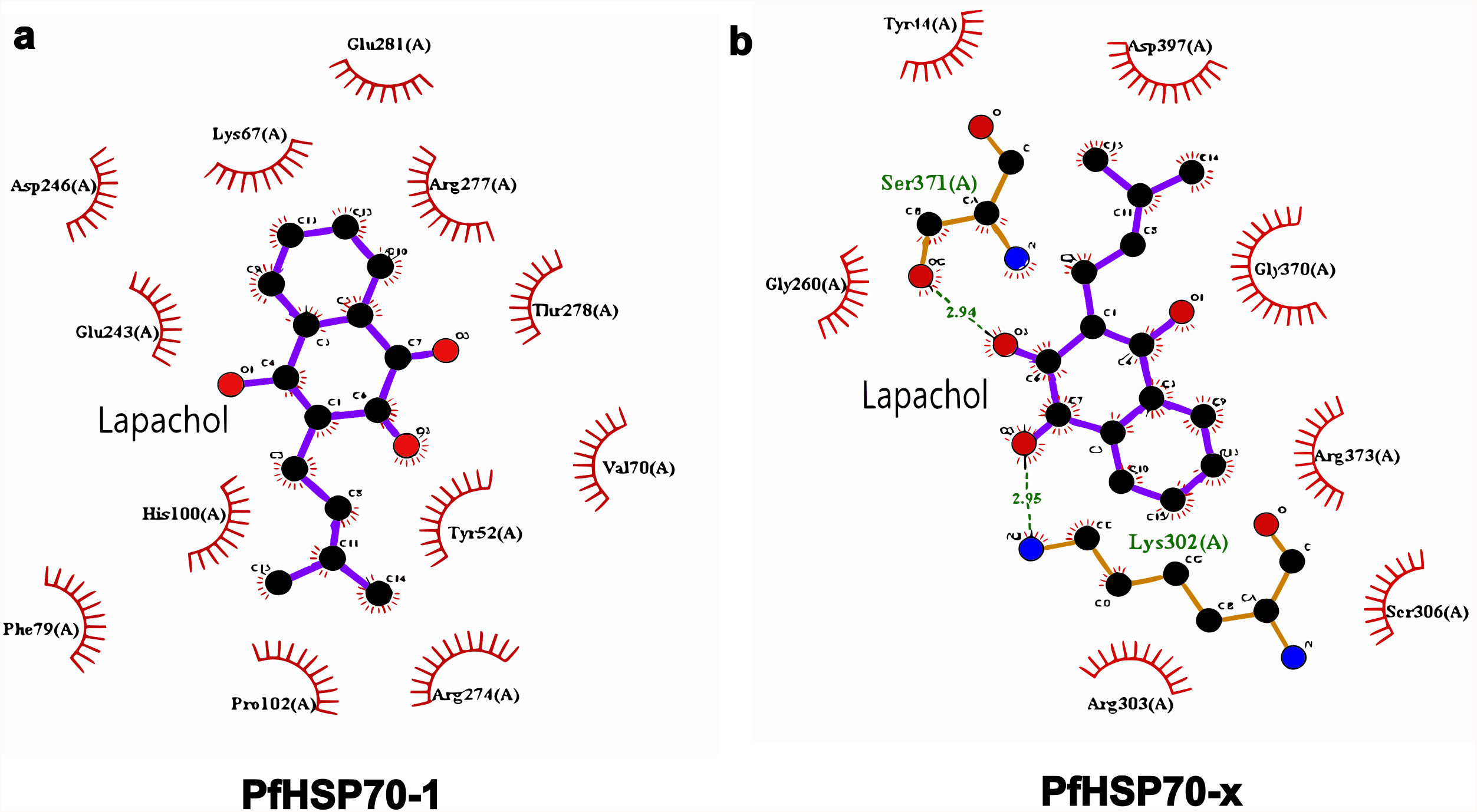
Ligplot analysis of selected complexesof lapachol with NBDs of. (a) PfHsp70-1(b) PfHSP70-x. Lapachol is shown in the center(ball and stick representation). Amino acids of PfHSP70s involved in non-covalent interactions like hydrophobic/ Van der Waals are depicted as stoked arcs. PfHSP70 residues involved in hydrogen bonding (green dotted lines) are shown as balls and sticks.

**Table 2:** List of interacting residues from PfHSP70 homologs. Amino acids from PfHSP70-1 and PfHSP70-x involved in lapachol interaction via hydrogen bonding or other non-covalent interactions like hydrophobic/ Van der Waals are listed in the table. Information was derived from Ligplot+ analysis.

| **PfHSP70 homolog** | **Lapachol binding residues** |
| --- | --- |
| PfHSP70-x | Y44, G231, G232, G260, K302, R303, S306, G370, S371, R373, D397, |
| PfHSP70-1 | Y52, K67, V70, F79, H100, P102, E243, D246, R274, R277, T278, E281 |

**Fig 5.**
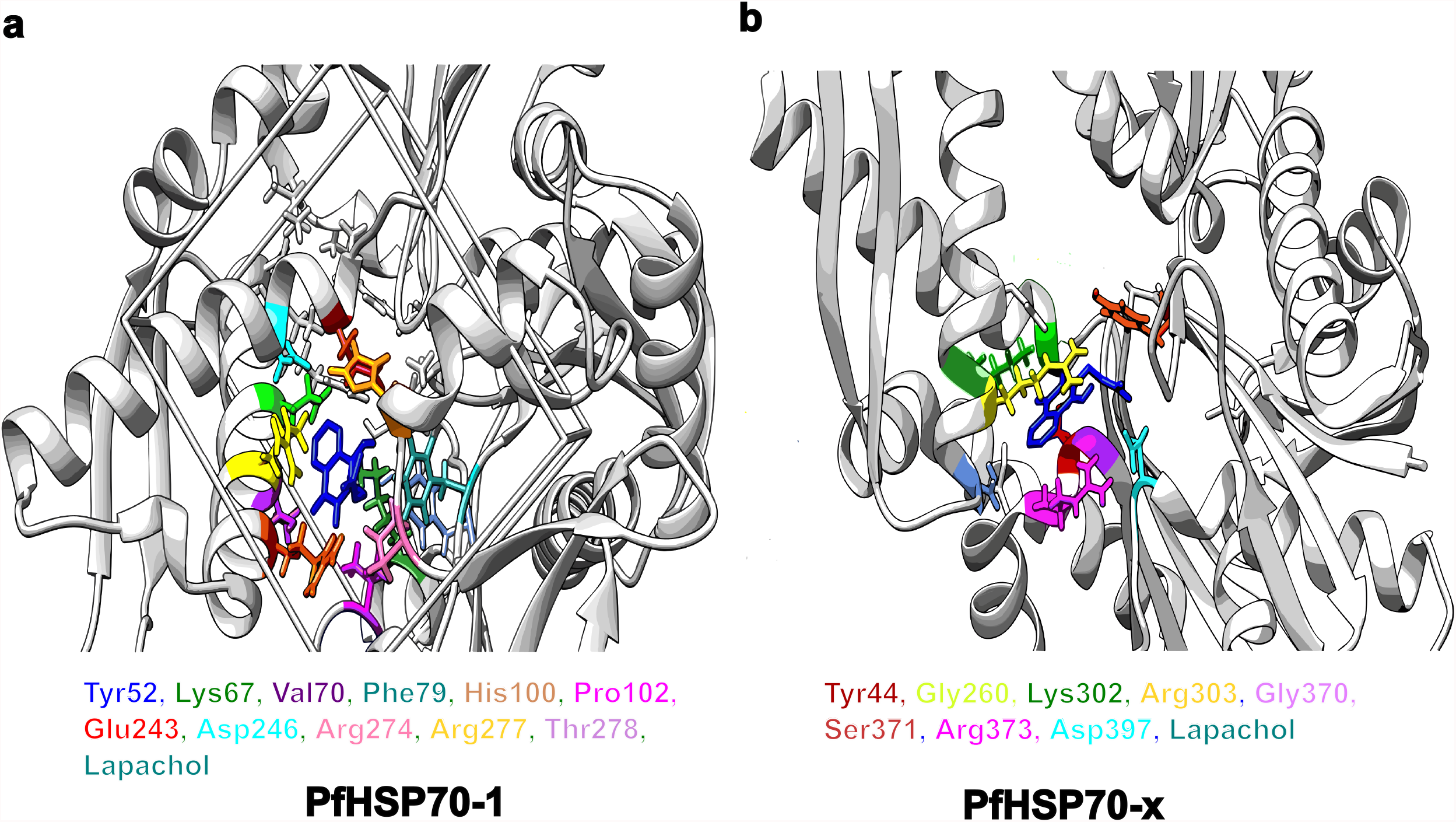
Cartoon representation highlighting lapachol binding pockets of PfHSP70 homologs. a. PfHSP70-1 and b. PfHSP70-x. Ribbon diagram of proteins is shown is grey. Amino acid residues of PfHSP70s involved in lapachol (dark blue) binding are shown as colored sticks. Side panels on each figure show names of highlighted residues in their respective colors on the cartoon.

## Conclusion

PfHsp70s are an important family of proteinsnecessary for parasite survival inside its two different hosts. A constant effort is being made to develop newer inhibitor molecules that can specifically target parasite Hsp70s. Lapachol was found to inhibit the ATPase activity of the exported PfHSP70-x in a concentration dependent manner whereas no significant effect of this molecule was seen on the nuclear/ cytoplasmic PfHSP70-1. In this study, we have docked lapachol on the three-dimensional structures of PfHSP70-1 and PfHSP70-x in an attempt to understand this differential inhibition of PfHSP70s by lapachol. Our results highlight a significant difference in the affinity of lapachol for these molecules as depicted by the binding energies of the obtained complexes(PfHSP70-x: -8.774; PfHSP70-1: -7.355). Analysis of these complexes further sheds light on the stark differences in the amino acid residues and types of interactions that PfHSP70-1 and PfHSP70-x perform with lapachol. Understanding these molecular interactions in detail may serve as the groundwork for design and development of new quinoneinhibitors against specific PfHSP70 homologs.

## Abbreviations

Pf: Plasmodium falciparum
HSP: Heat shock protein
SBD: Substrate binding domain
NBD: Nucleotide binding domain
ATP: Adenosine tri phosphate
ADP: Adenosine di phosphate

## Figure and table legends

**Fig S1: *Structure validation***. Ramachandran plotand ERRAT analysis of three-dimensional structures of PfHSP70-1 obtained from a. AlfaFold and b. SwissModel. The overall quality factors obtained from ERRAT for each model are mentioned below the plots.

## Acknowledgments

CN and VU were receiving scholarship from DBT, Government of India. SKis a DBT-SRF. The laboratories of PCM and RH are funded by UGC-DAE, and were earlier funded by DBT, RUSA and DST, Government of India.

## Declaration

The authors have no relevant financial or non-financial interests to disclose.

The authors declare that there are no conflicts of interest.

All authors read and approved the final manuscript.

